# Global and local redistribution of somatic mutations enable the prediction of functional XPD mutations in bladder cancer

**DOI:** 10.1101/2022.01.21.477237

**Authors:** Jayne A. Barbour, Tong Ou, Hu Fang, Noel C. Yue, Xiaoqiang Zhu, Michelle W. Wong-Brown, Haocheng Yang, Yuen T. Wong, Nikola A. Bowden, Song Wu, Jason W. H. Wong

**Affiliations:** School of Biomedical Sciences, Li Ka Shing Faculty of Medicine, The University of Hong Kong, Hong Kong Special Administrative Region, China; Urology Institute of Shenzhen University, The Third Affiliated Hospital of Shenzhen University, Shenzhen University, Shenzhen, China; Institute of Biomedical Data, South China Hospital, Medical School, Shenzhen University, Shenzhen, China; Centre for Drug Repurposing and Medicines Research, University of Newcastle, NSW, Australia; Hunter Medical Research Institute, Newcastle, NSW, Australia; Adult Cancer Program, Lowy Cancer Research Centre, UNSW Sydney, NSW, Australia; Department of Urology, South China Hospital, Medical School, Shenzhen University, Shenzhen, China; Centre for Oncology and Immunology, Hong Kong Science Park, Hong Kong SAR, China; Centre for PanorOmic Sciences, The University of Hong Kong, Pokfulam, Hong Kong Special Administrative Region, China

## Abstract

Xeroderma pigmentosum group D (XPD) is a DNA helicase with critical functions in transcription initiation and nucleotide excision repair. Missense mutations in XPD are putative drivers in around 10% of bladder cancers (BLCA), but the associated mutational process remains poorly understood. Here, we examine the somatic mutational landscape of XPD wild-type (n=343) and mutant (n=39) BLCA whole genomes. The genome-wide distribution of somatic mutations is significantly altered in XPD mutants, affecting both APOBEC and non-APOBEC associated mutational processes. Specifically, XPD mutants are enriched in T[C>T]N mutations (SBS2) with altered correlation with replication timing. At a locoregional genomic level, mutant XPD BLCA had striking T>G mutation hotspots at CTCF-cohesin binding sites (CBS) with evidence linking XPD to genomic uracil repair. Leveraging differential distribution of somatic mutations, we developed a machine-learning model for predicting pathogenic XPD mutations, which we validated in an independent TCGA cohort with 100% accuracy. Our model enabled the discovery of missed XPD mutation calls and uncovered pathogenic non-hotspot XPD mutations in bladder cancer. Our study reveals how XPD mutations redistribute somatic mutations in cancer genomes and provides a genome sequencing approach to differentiate driver and passenger XPD mutations.

## Introduction

Xeroderma pigmentosum group D (XPD), encoded by *ERCC2*, is a 5’-3’ ATP-dependent DNA helicase that is a component of the Transcription Factor II H (TFIIH) protein complex. TFIIH plays essential roles in transcription initiation through its interaction with RNA polymerase II (POLR2A) and nucleotide excision repair (NER) when recruited to damaged lesions. Compound heterozygous mutations in XPD can cause the recessive genetic disorders xeroderma pigmentosum (XP), Cockayne syndrome (CS) and trichothiodystrophy (TTD) which typically present with UV light sensitivity due to deficiencies in NER function [1]. These compound heterozygous mutations include the complete loss of function in one allele and a less deleterious point mutation in the other. The location of the XPD point mutations can vary but have been shown to affect the XPD’s helicase activity, stability and interactions with other TFIIH proteins [2, 3]. Somatic missense mutations in XPD are also putative drivers in cancers, with ∼10% of bladder cancer (BLCA) samples harbouring these alterations [4, 5]. XPD BLCA mutations do not overlap those underlying genetic disorders, but are commonly found in the helicase domains of XPD. XPD mutant BLCA are sensitive to cisplatin therapy, indicating a reduced capacity for repair of cisplatin adduct DNA lesions, implying a deficiency in NER of these samples [4, 6, 7].

Somatic mutation formation varies across the cancer genome in terms of mutation profiles and regional mutation densities. The type of single nucleotide variant (SNV, referred to as mutation) in the trinucleotide context forms specific single base substitution (SBS) mutational spectra or signatures which reflect the mutational process of the sample [8, 9]. Genomic mutation density can be highly varied across the genome, correlating strongly with various epigenetic marks such as chromatin accessibility [10], histone modifications [11, 12], transcription factor binding [13, 14] and cytosine methylation [15]. Regions of the genome with exceptionally high mutation densities are considered ‘mutational hotspots’. One such hotspot is CCCTC-binding factor (CTCF)-cohesin binding sites (CBS) of which there have been several reports of strongly elevated somatic mutation densities [16–21]. CTCF is a DNA binding protein that acts as a transcriptional repressor when bound to DNA alone and as an architectural protein when CTCF proteins bound at distal sites dimerise and interact with the cohesin complex to form DNA loops [22]. CBS hotspots have previously been linked to specific mutational signatures cosmic SBS7 [19] and SBS17 [16, 20]. SBS7 is an ultra-violet (UV) associated mutational pattern found in skin cancer and SBS17 is found in gastrointestinal cancers, particularly ESAD, and is characterised by T>G mutations [23]. SBS17 can be induced with treatment of 5-fluorouracil (5-FU) [24], a drug which increases uracil incorporation into DNA [25, 26].

XPD mutant BLCA has previously been associated with the enrichment of the mutational signature SBS5 [27]. However, how mutant XPD causes this mutational signature remains unknown. The other major mutational process occurring in BLCA can be attributed to the nucleic acid editing enzyme APOBEC [28]. APOBEC is a cytosine deaminase that deaminates cytosine to uracil, causing C>T mutations targeted at viral RNA but can also affect host DNA [28]. How XPD mutations alter processes of APOBEC mutagenesis has not been examined. To gain an insight into the XPD mutant-driven mutational process, we compared the mutation distribution of XPD wild-type and mutant BLCA across a range of genetic and epigenetic features. We found that XPD somatic mutations alter the mutational landscape of a range of mutational processes, and we demonstrate that this can be used to differentiate driver and passenger XPD mutations.

## Results

### Variable Contribution of APOBEC Associated and Other Mutations in a Cohort of 392 BLCA Samples

XPD mutations have been linked to a specific mutational signature in BLCA [27], but the genome-wide distribution of mutations associated with XPD mutants is unknown. To investigate this, we utilised the GE cohort of WGS BLCA and characterised samples that harboured putative XPD driver mutations. Out of a total of 392 samples, 39 were characterised as XPD mutant and 343 were characterised as WT due to the complete absence of protein altering XPD mutations. A further 10 samples were excluded from other analyses as they harboured a non-recurrent, protein altering mutation in XPD and we therefore could not confidently assign as either XPD mutant or WT.

To ensure that the phenotype responsible for these 39 samples is due to XPD, we compared the proportion of XPD mutant and WT samples with protein altering mutations in cancer drivers. We found that there are no other oncogene beside XPD is exclusively found in the XPD mutant samples (Figure S1A). Previous studies have found the presence of strong APOBEC mutational signatures in many BLCA samples [28] and signature 5 (SBS5) specifically in XPD mutant BLCA samples, but the analysis was restricted to exomes [27]. Since APOBEC related mutations are frequent in BLCA and have a distinct mutational process, we first assessed the contribution of APOBEC and non-APOBEC related processes (Other) in XPD mutant and non-mutant BLCA (Figure 1A). The spectrum of APOBEC and Other mutations overall showed a similar trend but with some differences in frequency at specific trinuleotide contexts (Figure 1B). We found that, as expected, the cosine similarity of APOBEC mutations was more similar to SBS2 and SBS13 than for all mutations, while other mutations were more similar to SBS5 than all mutations (Figure 1C).

**Figure 1.**
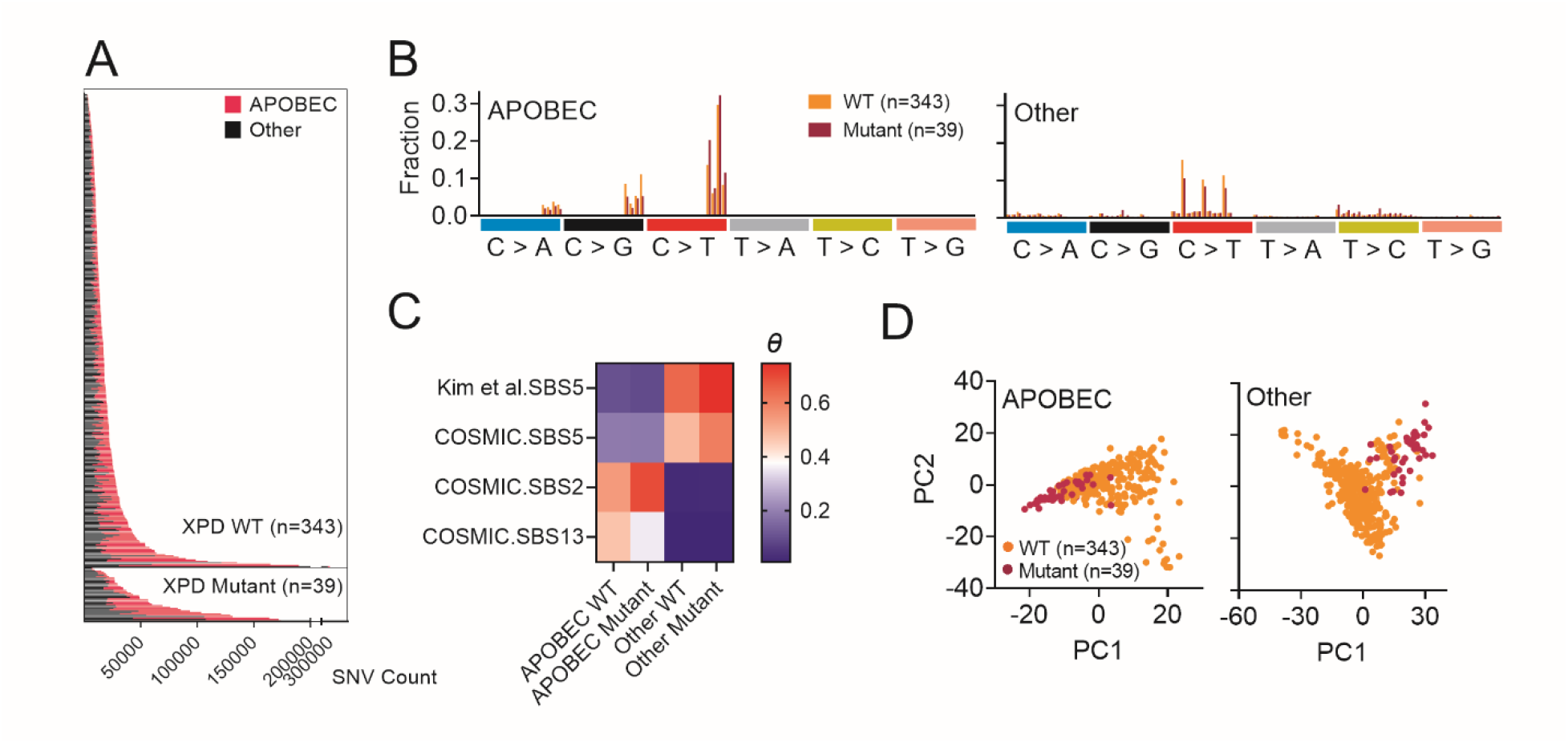
Contribution of APOBEC and Other mutations in XPD mutant and WT BLCA. (A) Total number of mutations attributed to T[C>D]N (APOBEC) or not T[C>D]N (Other) in GE WGS BLCA cohort arranged by genotype and total mutation number. (B) Trinucleotide mutational spectra of APOBEC and Other mutations for GE WT (orange) and XPD mutant (pink) BLCA samples. (C) Heat map of cosine similarities to mutational signatures in BLCA where theta = 1 is most similar. COSMIC signatures 2, 5 and 13 are from COSMIC, the other signature, TCGA.130.DFCI.MSK.50.signature5 is from the supplementary material from [27] (D) Principle component (PC) analysis plots representing PC1 and PC2 of observed-expected mutation density ratios were calculated across each 1 mb window of hg38 for all SNVs, APOBEC SNVs and Other SNVs

Other SNVs also contained C>T mutations at CpGs which is a common mutation caused by deamination of methylcytosine that is displayed in all cancer types. We next investigated the distribution of APOBEC and Other mutations in 1 Mb windows across the genome. We found that the genome-wide distribution of All, Other and APOBEC mutations differ between WT and XPD mutant samples (Figure 1D).

### XPD Mutant Samples Display Altered Genomic Distribution of Somatic Mutations

Mutations in most mismatch repair proficient cancers show variation in mutation burden in relation to replication timing, chromatin accessibility and gene expression [10]. To explore the relationship between mutation density and these epigenomic features further, mutation densities for APOBEC and Other mutations were calculated with respect to gene bodies and replication time in XPD mutant and WT BLCA. Expressing mutation density as the observed-expected mutation ratio (see Methods), we found that the distribution of APOBEC and Other mutations was significantly higher in all genic regions and lower in intergenic regions in XPD mutant samples compared with WT (Figure 2A). We found an increase in the burden of APOBEC related mutations in 5’UTR relative to what is expected by chance (observed-expected ratio > 1) in both XPD mutant and WT groups (Figure 2A). The burden of APOBEC related mutations was increased in 5’UTR in both XPD mutant and WT groups (Figure 2A), consistent with previous findings that APOBEC causes mutation clusters around the start of active genes which could be from ssDNA exposure as transcription commences [43]. A linear regression between Other mutation densities and replication time showed that WT samples had a slope of -0.01382 compared with -0.002548 for mutant (Figure 2B), with significantly decreased and increased burdens of mutations in mutant samples compared with WT in late and early replicating regions respectively (q=0.000086 and q=<0.000001, Student’s t-test with multiple testing correction, Figure S2A). APOBEC mutations had a striking trend of being negatively associated with replication time in WT samples (slope = -0.007405) but positively associated with replication time in XPD mutant samples (slope = 0.006413) (Figure 2B, Figure S2A). The observed relationship between replication time and XPD mutant BLCA is consistent across all mutation types (Figure S2B). The differential burden of Other mutations between mutant and WT samples in gene bodies and over the replication time landscape suggested that transcriptionally active or open chromatin plays a role in the distribution of XPD related mutagenesis. We next examined the effect that transcriptional activity has on mutagenesis in the BLCA genomes. We find that XPD mutant samples have increased genic mutation burden for both Other and APOBEC mutations, compared to WT specifically at expressed genes (Figure 2C). This was particularly pronounced immediately before the transcriptional start site (TSS) (Figure 2C and S2C).

**Figure 2.**
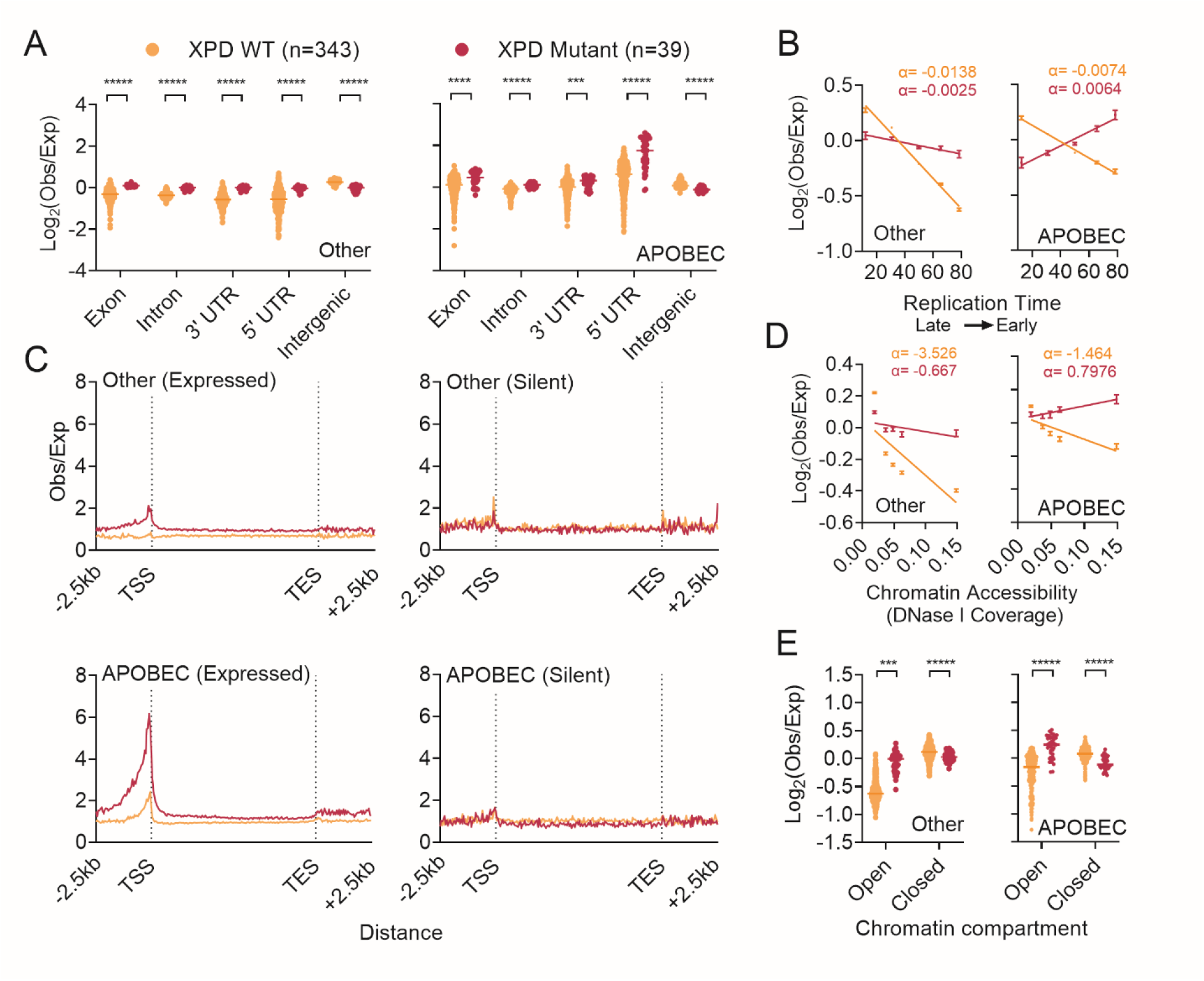
Genome-wide distribution of APOBEC and Other mutations in XPD mutant and WT BLCA. (A) Mutation densities as observed-expected ratios (obs/exp) in exons, and 3’ and 5’ untranslated regions (UTR), introns or not in any of these regions (intergenic) in WT samples in orange and XPD Mutant samples (Mutant) in pink with Other SNVs displayed on the left and APOBEC SNVs displayed on the right. (B) Mutation densities as obs/exp for 5 genomic bins organised by replication time. BrDU immunoprecipitation coverage was used for binning where a higher number indicates earlier replication time. Plots and error bars represent mean and standard deviation of different samples and the line represents a linear regression model between mutation densities and the mean replication time for each of the bins. (C) Observed-expected mutation density ratio profile plots Other SNVs (left) and APOBEC SNVs (right) across gene body of genes expressed in bladder tissue (Expressed Genes) or genes not expressed in bladder tissue (Silent Genes). (TSS = transcriptional start site, TES = transcriptional end site). The gene body was organised into 150 bins and the region 2.5 Kb up or downstream of the TSS or TES was organised into 50 bins. (D) Mutation densities as obs/exp for 5 genomic bins organised by chromatin accessibility measured by DNase hypersensitive coverage where a higher number is more accessible. Plots and error bars represent mean and standard deviation of different samples and the line represents a linear regression model between mutation densities and the mean DHS coverage for each of the bins. (E) Observed/expected mutation density ratios for genomic regions annotated in normal bladder to be either a chromatin A compartment (open) or chromatin B compartment (closed) published [38]. *** q < 0.0001, ***** q < 0.000001, Student’s t-test with multiple testing correction.

To look more generally at active and inactive chromatin genome-wide, we next used DNase hypersensitivity (DHS) to compare the burden of APOBEC and Other mutations between XPD mutant and WT samples. A regression was performed between DHS coverage and somatic mutation densities across 5 bins and we found a marked difference in the relationship between XPD mutant and WT cancer. XPD mutant and WT samples had slopes of -3.526 and -0.6667 respectively for Other SNVs and -1.464 and 0.7976 for APOBEC SNVs. For APOBEC SNVs, there were significantly more mutations in XPD mutants compared with WT in most accessible DHS regions and significantly less in less accessible regions (q<0.000001, Student’s t-test with multiple testing correction, Figure 2D). We also found significantly increased and decreased burden of Other mutations in XPD mutant samples in open compartments and closed compartments respectively compared with WT (q< 0.000001, q=0.000024, Student’s t-test with multiple testing correction, Figure 2E). Exclusion of CBS and genes did not affect the observed assocations (Figure S2D, S2E). This provides strong evidence that XPD protects accessible chromatin from mutagenesis.

### XPD mutant cancers are enriched in APOBEC induced cytosine deamination associated mutations

APOBEC mutational signatures can be further divided into SBS2 and SBS13, where SBS2 is dominated by T[C>T]N and SBS13 is dominated by T[C>R]N (where R is A or G). Earlier, we showed that XPD mutant BLCA have higher cosine similarity to SBS2 and lower cosine similarity to SBS13 compared with WT BLCA (Figure 1C). The differences in the distribution of APOBEC mutations across the replication time landscape were also particularly striking with WT having a slope of -0.007405 while XPD mutants having a slope of 0.006413 (Figure 2B). So we next wanted to explore APOBEC related mutagenesis in more detail and separated SBS2 and SBS13 for futher analysis.

We found that XPD mutant BLCA have a higher contribution from SBS2 and lower contribution from SBS13 than WT (Figure 3A). Moreover, the fraction of APOBEC mutations being T[C>T]N and T[C>R]N were higher and lower respectively than WT (Figure 3B). APOBEC associated T[C>T]N is known to be caused by mutations caused by unrepaired U>G mismatches that result from APOBEC induced deamination of cytosine to uracil [28]. APOBEC overexpressing yeast acquire C>D somatic mutations whereas uracil-DNA glycolase (*UNG*) mutant, APOBEC expressing yeast exclusively acquire C>T somatic mutations [44]. A recent study also found that human cancer cell lines with strong endogenous APOBEC signatures were enriched in SBS2 when they are *UNG* deficient [30].

**Figure 3.**
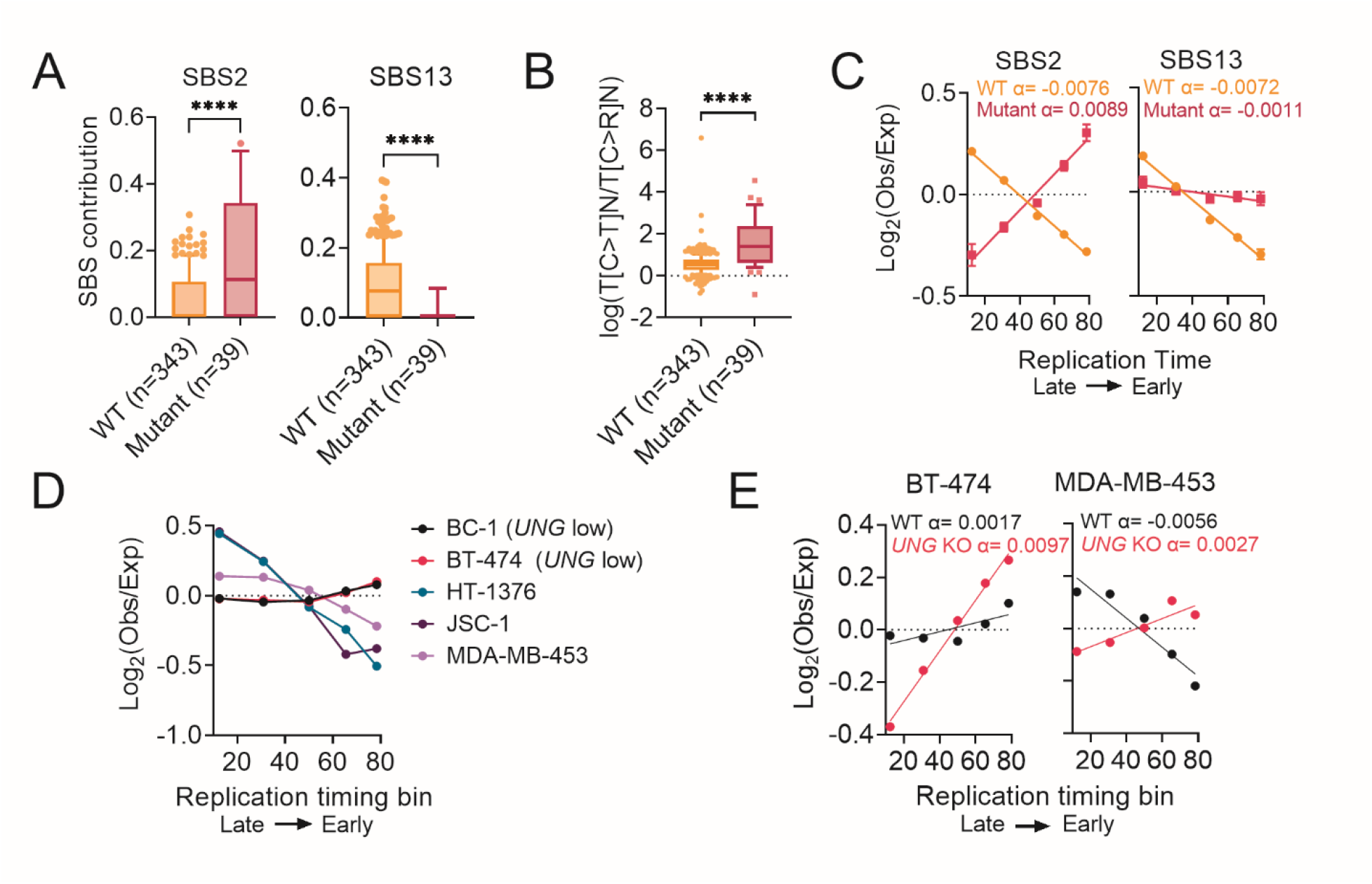
APOBEC mutagenesis in WT and XPD mutant cancer. (A) The weighted signature fraction contribution of SBS2 and SBS13 from deconstructSigs for XPD mutant and WT BLCA. (B) The log ratio of APOBEC mutations that can be attributed to T[C>T]N or T[C>R]N where R is A or G. (C) Mutation densities for T[C>T]N (SBS2) and T[C>R]N (SBS13) mutations as obs/exp for 5 genomic bins organised by replication time for XPD mutant and WT BLCA. (D) Mutation densities as obs/exp for 5 genomic bins organised by replication time for the following cell lines that contain endogenous APOBEC mutational signatures – BC-1, BT-474, HT-1376, JSC-1 and MDA-MB-453. (E) Mutation densities as obs/exp for 5 genomic bins organised by replication time for WT and *UNG* knockout CRISPR in BT-474 and MDA-MB-453 cells. Mutation data for cell lines is from [30]. **** is p<0.00001 by unpaired t-tests

We next examined the relationship between APOBEC associated mutations and replication time. We found that T[C>T]N mutations had a positive relationship with replication time in XPD mutant samples (slope= 0.008926) but a negative relationship with replication time in WT samples (slope = -0.007569)(Figure 3C). T[C>R]N mutations on the other hand, display a less prominent difference in its relationship with replication time between WT and XPD mutant with slopes of -0.007186 and -0.001147 respectively (Figure 3C).

We reanalysed mutation patterns in cancer cell lines with APOBEC signatures [31] for comparison to WT and XPD mutant BLCA in order to dissect processes of APOBEC mutagenesis. BC-1 and BT-747 cells, which have low *UNG* expression, both displayed a positive relationship between T[C>T]N mutations and replication time, while the other *UNG* proficient cell lines showed a strong negative trend (Figure 3D), corresponding the XPD mutant and WT profiles, respectively. The study also profiled somatic mutations accumulated by cells with knock-out (KO) of genes related to the APOBEC pathway including *UNG*. Both BT-474 and MDA-MB-453 KO cell lines displayed redistribution of T[C>T]N mutations towards a positive association with replication time (Figure 3E). Thus, APOBEC associated mutations in XPD mutant BLCA share characteristics of mutations associated cytosine deamination induced genomic uracil in *UNG* deficient cells.

### XPD Mutant Cancers Display Strong Mutation Hotspots in CTCF-Cohesin Binding Sites

DHS are associated with *cis*-regulatory elements, including promoters, enhancers and CTCF-cohesin binding sites (CBS). As mutations in XPD mutants showed increased mutation density at DHS regions (Figure 2D), we examined the different classes of cis-regulatory regions and found striking mutation hotpots in XPD mutants at CBS, with only minor increase in mutation rate at promoters or enhancers (Figure 4A). We similarly observed CBS hotspots in a total of 7 XPD mutant BLCA samples from three other studies [31–33] (Figure S3A) as well as 3 XPD mutant liver cancer samples from PCAWG [31] (Figure S3B).

**Figure 4.**
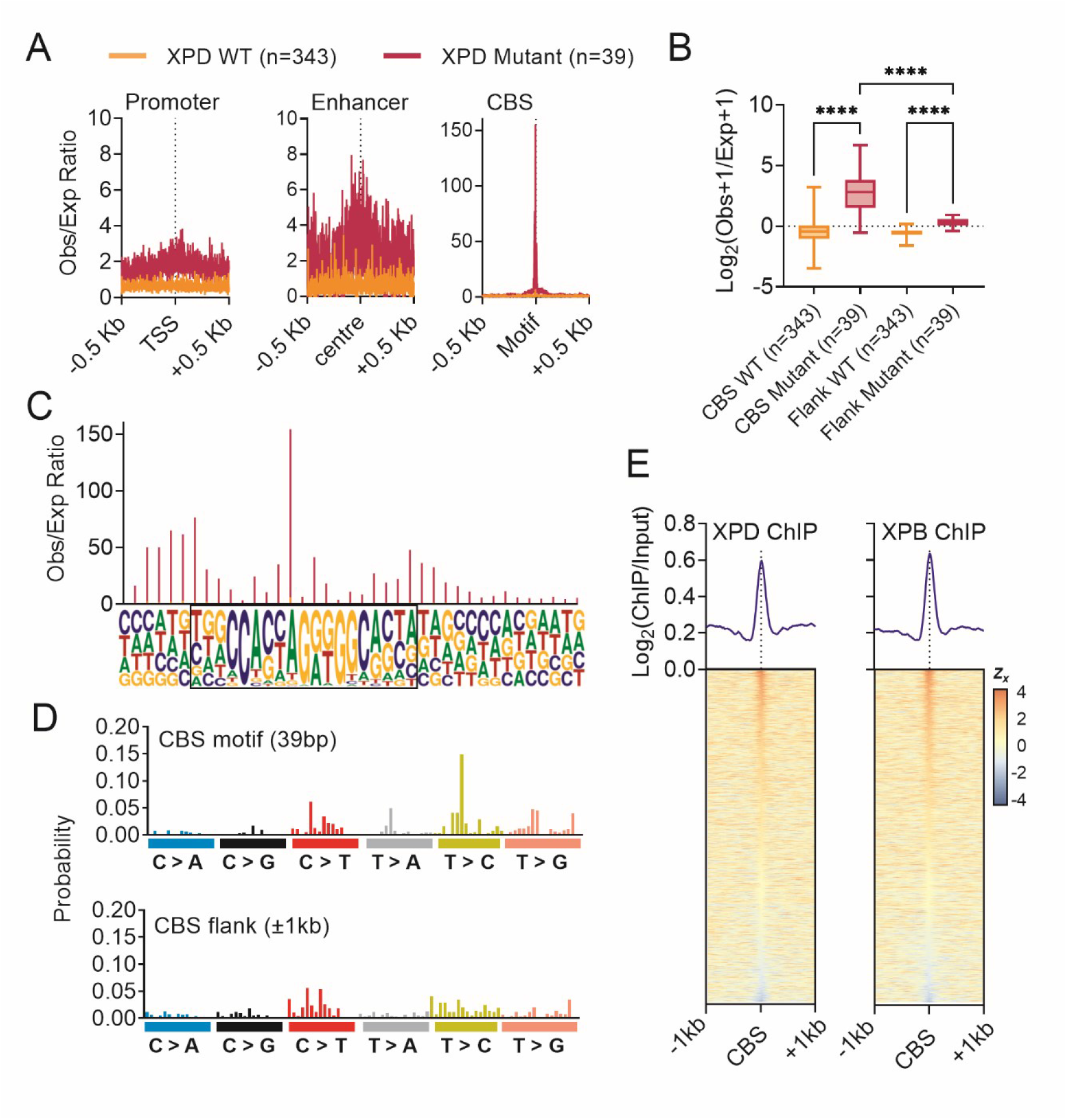
Mutation densities at DNase hypersensitive regions in XPD mutant and WT BLCA. (A) Profile plots of mutation densities as observed-expected ratios (obs/exp) for Other Mutations in regions annotated in normal bladder as the promoter, enhancer or CTCF-Cohesin binding site (CBS) 0.5 Kb up or downstream of the TSS, centre and motif, respectively. (B) Mutation densities (obs/exp) of CBS motif and +/-1 Kb flanking regions for WT and XPD mutant GE BLCA samples **** q < 0.0001, Student’s t-test with multiple testing correction. (C) Observed/expected mutational profile of GE XPD mutant samples across the CBS motif. (D) Trinucleotide mutation frequencies of bladder XPD mutations in CBS and flanking regions. This was normalised using manual normalisation in deconstructSigs to account for the trinucleotide composition of the regions. ***** q < 0.000001, Student’s t-test with multiple testing correction. (E) Profile and heat maps of coverage of XPD and XPB ChIP-seq across CBS. Data accessed from (GEO:GSE44849).

While somatic mutation hotspots in CBS have previously been reported in UV associated skin cancers [19] and SBS17 associated gastrointestinal cancers [16, 18, 20], it is a striking and novel observation that XPD mutant BLCA also have CBS mutation hotspots. We also observed increased mutation densities in flanking regions in XPD mutant samples compared with WT which may reflect generally greater chromatin accessibility of the CBS flank and, but is substantially lower compared with the CBS itself (Figure 4B). APOBEC SNVs also displayed elevated mutation densities at CBS for XPD mutant but not WT BLCA (Figure S3C). However, these elevated mutation density profiles across the CTCF motif were not as striking in terms of observed expected ratios as Other SNVs (Figure 4A and S3C). Previous reports of CBS hotspots had found specific mutational patterns and signatures across the CBS motif [16, 19]. We observed that the CBS mutations in XPD mutants also have a similar distribution across the CTCF motif compared with CBS hotspots found in esophageal adenocarcinoma (ESAD) (Figure 4C, Figure S3D), but is different to melanoma (MELA) (Figure S3E). This potentially implicates a shared mechanism of CBS mutagenesis with ESAD. In terms of the type of the CBS specific trinucleotide mutational spectrum, there is strong enrichment for T>N mutations with the strongest enrichment being T>G which is absent from the CBS flank (Figure 4D). This is similar to gastroesophageal cancers with SBS17 where predominantly T>G and T>C mutations accumulate at CBS [16, 20].

### Effect of XPD Presence on Genomic Mutation Distribution

We next wanted to explore how XPD itself affects the development of mutations in WT and mutant cancers. Using previously published XPD ChIP-seq data [39], we examined XPD’s distribution around CBS. We found a strong enrichment of XPD at CBS, and as XPB is also enriched at CBS (Figure 4E), implying that these proteins are co-bound to CBS as part of the TFIIH complex. Genome-wide nucleotide excision repair activity mediated by TFIIH in response to cisplatin has previously profiled by XR-seq [45]. XPD mutant BLCA are sensitive to cisplatin suggesting they have a role in repairing this type of damage [5, 7]. Using the XR-seq data we correlated TFIIH repair activity and mutation burden in XPD mutant and WT BLCA. We found a strong inverse correlation between mutation densities and TFIIH repair in WT BLCA samples, whereas this trend was completely lost for XPD mutant (-0.09581 versus 0.02224, FigureS4B) with significantly higher obs/exp and lower obs/exp in WT compared with mutant for high and low TFIIH coverage regions respectively (q=0.000003 and q=0.000178, Student’s t-test with multiple testing correction, Figure S4B).

Elevated mutation burdens in high TFIIH repair regions in XPD mutant BLCA indicates that mutagenesis in these regions is likely caused by a loss of TFIIH mediated DNA repair. To further delineate this, we examined the mutational spectrum in high and low TFIIH regions of the genome. We rationalised that if mutant XPD is actively causing a specific sort of damage, then XPD mutant samples will have a specific mutational signature at high TFIIH regions. We found that the mutational spectrum was highly similar between low and high TFIIH regions in both XPD WT and mutant cancers (Figure S4C). Additionally, WT and XPD mutant cancers had a similar signature to each other in high TFIIH regions (Figure S4C). Together these results illustrate that the presence of TFIIH repair affects the mutation burden but not mutation types which is consistent with a loss of DNA repair.

### Genomic uracil accumulates at CTCF-cohesin binding sites

Earlier we found that XPD mutant may be associated with dysfunctional repair of genomic uracil that result from APOBEC associated cytosine deamination in BLCA. To determine if genomic uracil might also be associated with CBS mutagenesis, we took advantage of previously published base pair resolution sequencing data of genomic uracil in *UNG* KO cells [41]. Since the data is base pair resolution, uracil from incorporation and deamination can be identified based on whether the reference base is A/T or C/G respectively. We found that there is strong enrichment of genomic uracil at CBS (Figure 5A). The distribution of uracil across the motif is asymmetric, mirroring XPD mutant CBS mutation hotspots (Figure 5B). Treatment with pemetrexed (PMX), which increases uracil misincorporation, results in an even greater enrichment of uracil at CBS (Figure 5A). Analysis of the trinucleotide context of uracil misincorporation show that the frequency of uracil incorporation sites was most enriched in TTT followed by CTT (Figure 5C). This resemebles trinucleotides most strongly mutated in the CBS motif in XPD mutant cancers and is also the most prevalent trinucleotides context in SBS17, where CBS hotspots are also observed [16, 17, 20]. These finding suggest that the CBS mutational hotspots in XPD mutants and cancers with SBS17 maybe be associated with the repair and misincorporation of uracil in DNA.

**Figure 5.**
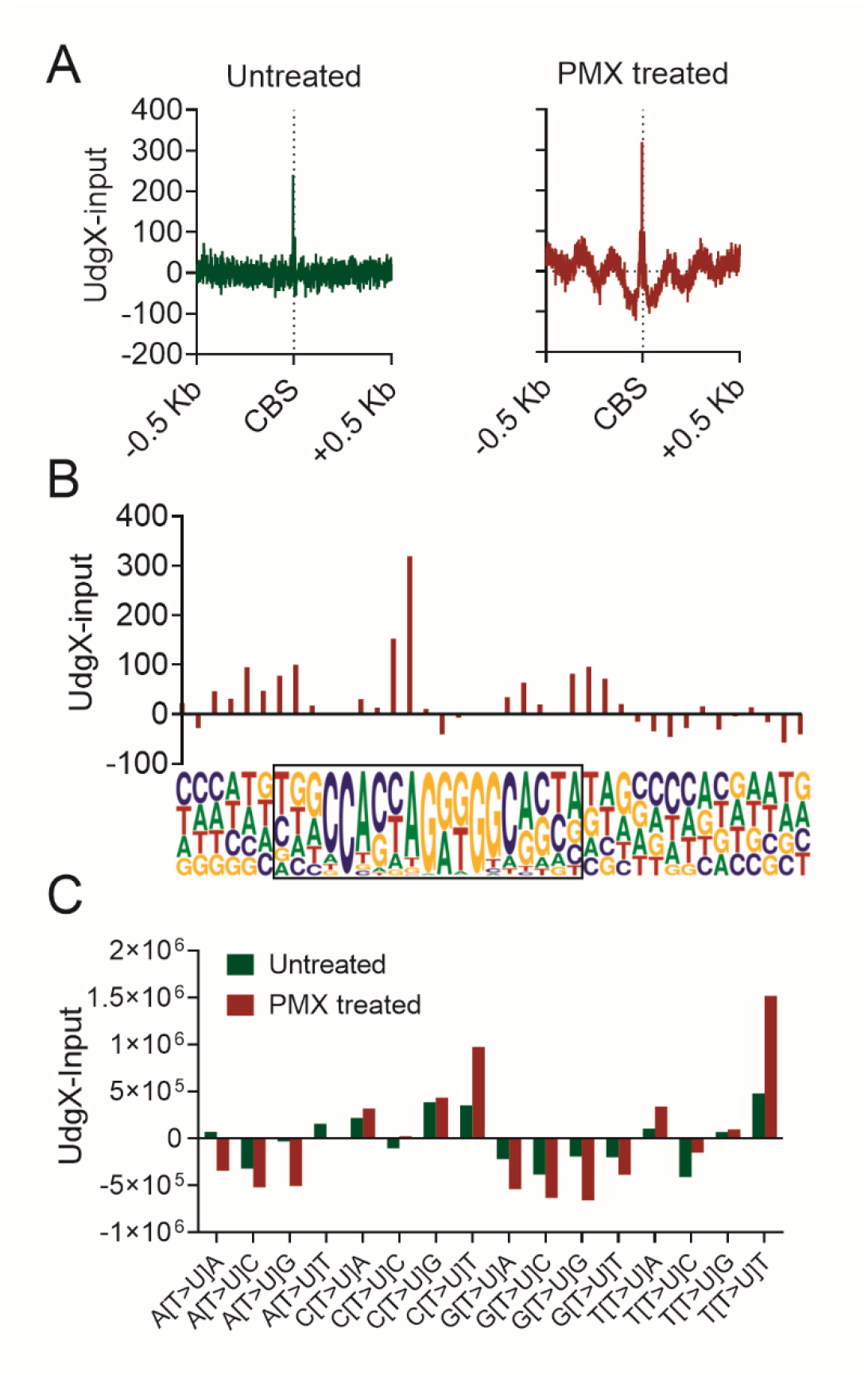
Genomic uracil distribution at CTCF-Cohesin binding sites. (A) Profiles of UdgX sequencing data (SRA:SRP319102) as input (inp) subtracted from experiment (exp) across CBS with 0.5 Kb flank up-(+) and down-(-) stream. (B) UdgX sequencing exp-inp across each base of the CTCF motif of CBS. (C) Frequency of single base sites of uracil incorporation in the trinucleotide context as UdgX sequencing exp-input. T>U indicates a uracil was detected where the reference base is a thymine

### Genome Wide Distribution of Somatic Mutations Predicts XPD Mutation Status in BLCA

Cancer patients with tumours harbouring XPD mutations have favourable responses to cisplatin [4, 6]. XPD is included in most targeted gene sequencing panels for molecular diagnosis of cancer, however, the pathogenicity of XPD mutations is not always clear as they often occur outside of known hotspots [6]. We therefore wanted to test if it is possible to predict a sample’s XPD mutation status based on its genome-wide distribution of somatic mutations with the idea that WGS data could be used to verify the pathogenicity of uncertain XPD mutations. To this end, we generated SVM models to classify whether a sample is XPD mutant based on the local and global distribution of somatic mutations.

First, using leave-one-out-cross validation on the GE BLCA cohort with well-defined XPD mutation status (WT = 343, mutant = 39), the SVM models achieved 98.69% accuracy (sensitivity:97.22%, specificity 98.84%, Figure 6A). One of the WT samples that was misclassified was predicted to have the highest probability of being a mutant. We manually examined the unfiltered vcf file for mutations in XPD in this sample, and found that the sample in fact contains a N238T mutation at relative low variant allele frequency (4/84 reads) which was filtered by the variant caller. N238 is a known mutation hotspot in bladder cancer, thus our classifier is also able to identify functional XPD mutations that are missed by variant callers. We further evaluated the SVM model trained on the GE cohort using an independent TCGA WGS BLCA cohort (WT = 19, mutant = 4), achieving 100% accuracy in classifying both WT and mutant samples (Figure 6B). To determine the relative importance of the mutational features in predicting XPD mutation status, a sensitivity analysis was performed on the SVM model. This found that the observed/expected ratio of Other mutations at CBS was by far most important, followed by the observed/expected ratio of APOBEC mutations at CBS (Figure 6C), supporting our observation at CBS hotspots are highly distinctive in XPD mutant cancers.

**Figure 6.**
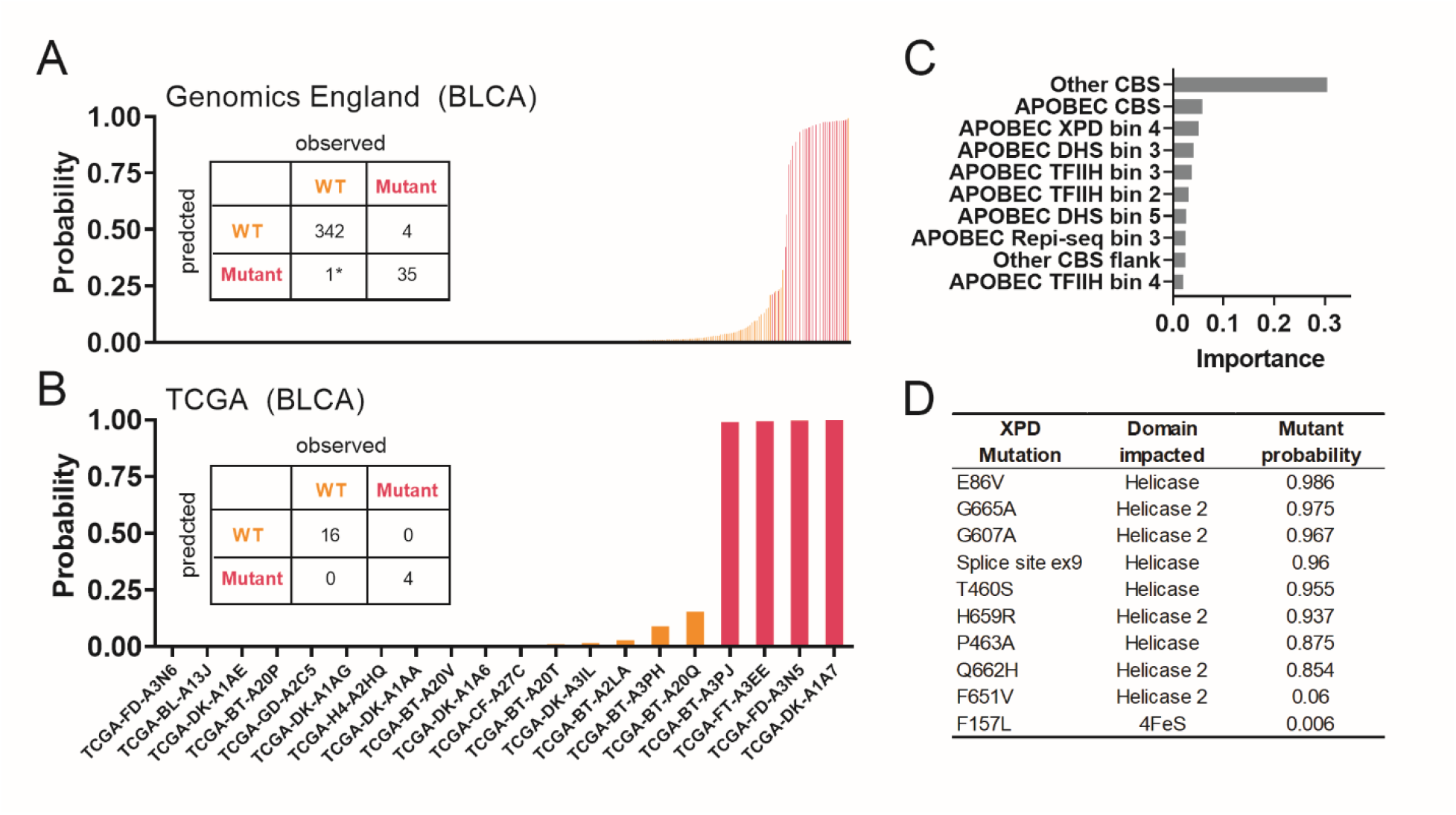
Support vector machine model predicts XPD Mutation Status based of genome-wide distribution of somatic mutations. (A) Classification of samples as WT (orange) or XPD mutant (pink) from SVM and associated confusion matrix showing the accuracy of predictions by leave-one-out cross validation of the GE cohort. (B) Classification of samples as WT (orange) or XPD mutant (pink) from SVM and associated confusion matrix showing the accuracy of predictions of PCAWG BLCA samples. (C) Sensitivity analysis showing importance of features as predictors in SVM model (D) Prediction from SVM when applied to 10 GE samples that had non-recurrent but protein altering mutations in XPD. Probability is the prediction from the model and Domain impacted refers to the protein domain of XPD that where the mutation is located.

Finally, we apply our SVM model to the Genomics England BLCA cohort which we had excluded from our earlier analysis as their XPD mutation status is uncertain due to the mutations not being located at known hotspots (n = 10). The SVM predicted all except two samples to be XPD mutant. The two samples predicted to be wild-type included one with a F157L mutations and one with a F651V (Figure 6D). F157L is outside of the helicase domains, while F651V is not within a conserved helicase motif, which may explain the lack of functional effect in these samples.

## Discussion

In this study, we show that XPD mutant cancers display distinctive genomic distribution of somatic mutations including mutation hotspots at CTCF-cohesin binding sites. We leverage this knowledge to build a SVM model that differentiates driver and passenger XPD mutations. This is particularly useful for mutations in XPD as pathogenic mutations appear as point mutations that are widely distributed across the protein. Pathogencity is currently inferred based on hotspot sites found across BLCA patients. However, this means that pathogenic mutations appearing at rarer sites can be missed. Our SVM model, we were not only able correctly classify XPD mutant samples with 99% accuracy, but we also identify one sample where the XPD mutation was filtered due to low variant allele frequency. Using our SVM model, we further identify 8 samples with non-recurrent functional XPD mutations (from over 120,000 cancer samples, including over 3,000 BLCA samples). This WGS approach to detect pathogenic XPD mutations may have clinical significance, as BLCA patients with these mutations are more sensitive to cisplatin treatment [5, 7].

While the biological mechanisms of this observation require further investigation, our results support a role for XPD mutants in compromised DNA repair. We observed an inverse relationship between TFIIH repair coverage and mutation densities in WT BLCA samples which is lost in mutant samples. This indicates that XPD is playing a protective role in the WT samples and is absent in mutants (Figure S4B). Further, the mutational signatures were highly similar in high and low TFIIH repair regions indicating that a similar mutational process is taking place where XPD is bound compared to where XPD is not bound (Figure S4C). The fact that the mutation spectrum is unchanged, but the mutation burden is increased is consistent with a loss of DNA repair. These results are also consistent with the clinical disease progression of patients with XP caused by XPD mutation where the incidence of skin cancer is greatly increased in sunlight exposed skin [46]. This supports the idea that mutant XPD protein does not directly generate DNA damage, but rather, is unable to repair damage arising from genomic insults such as UV-light.

Analysis of mutations from cell lines with endogenous APOBEC signatures and KO of APOBEC related genes revealed that the mutations of XPD mutant BLCA mirror that of *UNG* deficient cell lines (Figure 3D and 3E). *UNG* encodes an uracil excision enzyme (UDG2) therefore implicating a role for XPD in protecting CBS from genomic uracil. Interestingly, SBS17 which is also associated with mutation hotspots at CBS may also be caused by genomic uracil. SBS17 can be recapitulated in cell culture experiments by treatment with 5-FU and SBS17 mutations develops in breast and colorectal cancer tumours where patients were treated with 5-FU [24]. dUTP can be incorporated into DNA by mammalian polymerases at similar efficiencies to dTTP but the cell gets around this by keeping dUTP levels low. However, 5FU is a thymidylate synthase inhibitor which increases the dUTP/dTTP ratio in the cell leading to genomic uracil incorporation. Incorporation of dUTP can cause T>G mutations [47–49] which is the hallmark of SBS17. UDG2 does not distinguish between dUTP generated by cytosine deamination attributed to APOBEC activity or by erroneous misincorporation of dUTP from the free nucleotide pool such as those from 5-FU treatment [50]. Therefore, these seemingly unrelated mutational processes of APOBEC cytidine deamination in BLCA and uracil misincorporation in SBS17 may be related by uracil excision, whether it be by deamination or incorporation. Given that uracil misincorporation is not directly mutagenic, mutant XPD likely contributes to defective repair of abasic sites following the excision of uracil by UDG2. Further experiments will be required to fully elucidate the role of XPD in the repair of genomic uracil.

## Materials and Methods

### Sample Information

Somatic mutation calls from 392 Genomics England (GE) whole-genome sequenced (WGS) BLCA samples were accessed directly from the GE research environment. The Memorial Sloan Kettering (MSK) cohort from the AACR-GENIE database was used to find recurrent XPD mutations with recurrence being defined as a protein altering mutation in XPD found in one BLCA sample plus one other sample of any cancer type.

Of the 392 GE BLCA samples, 39 contained recurrent XPD mutations which we grouped as ‘XPD mutant’ and 343 samples had no protein altering mutations in XPD which we grouped as ‘WT’. A further 10 samples had protein altering mutations in XPD that were not recurrent. These 10 samples were excluded from analysis as we could not confidently assign as either WT or XPD mutant.

To analyse the proportion of oncogene protein altering mutations in WT and XPD mutant groups, we counted the number of samples in each group with either missense, nonsense or frameshift mutations in genes from the IntOGen database [29].

For SBS17 and SBS7 cancers we used C>T and T>G mutations from GE esophageal adenocarcinoma (ESAD) and all GE melanoma (MELA), respectively. This resulted in 828,349 C>T and 794,567 T>G mutations from 106 samples for ESAD and 363,284,245 mutations from 337 samples for MELA.

For analysis of APOBEC mutations in cell lines, data was accessed from [30]. This dataset included somatic mutation data from single cell clones and included the following cell lines with endogenous APOBEC mutational signatures – BC-1, BT-474, HT-1376, JSC-1 and MDA-MB-453. BT-474 and MDA-MB-453 cells also mutations from clones that were either WT and *UNG* (encoding UDG2) knockout by CRISPR. We pooled daughter mutations together by cell line.

We additionally utilised somatic mutations from Pan-cancer analysis of whole genomes (PCAWG) BLCA and liver cancer [31], a study of WGS urothelial bladder carcinomas [32] and a study of WGS neuroendocrine bladder cancer [33]. The PCAWG BLCA study had 4 XPD mutant and 19 WT samples. The urothelial bladder carcinoma study had 2 XPD mutant and 63 WT.The neuroendocrine BLCA had 1 XPD mutant and 5 WT samples. These 3 studies of BLCA were combined giving 7 XPD mutant and 87 WT. For liver cancer samples from PCAWG, there were 3 XPD mutant samples and 302 WT. These mutation calls were hg19 and hg19 annotations were used to generate these figures.

### Somatic mutations and simulation

SNV calls for hg38 were obtained directly from the GE research environment (RE). For BLCA, SNVs that were C>D (D represents A, G or T) at TCN context were defined as APOBEC whilst all SNVs not in this context were defined as ‘Other’ (Other). For ESAD, SNVs were classified as either C>T or T>G regardless of trinucleotide context. Simulations were used to establish the chance of genomic positions being mutated based on sequence context and mutation burden using SigProfilerSimulator [34]. For BLCA and ESAD, 100 simulations were performed and were merged and divided by 100 giving what we refer to as ‘expected’. For MELA, 10 simulations were used instead of 100 due to memory constraints.

### Calculation of Local Mutation Densities and Generation of Mutation Profiles across Genomic Sites

To calculate mutation densities at specific genomic regions, we counted the number of actual mutations (observed) and simulated mutations overlapping these regions using the tool ‘intersectBed’ [35]. Mutations of 100 simulations were merged for analysis and then divided by 100 to give an ‘expected’ value, and then local mutation density was expressed as the ratio of observed to expected mutations.

Genome-wide distribution of mutations was performed by calculating mutation densities as described above for 1 megabase (mb) windows of the human genome and then principal component analysis (PCA) was performed in R using prcomp() function with scaling and centering. To perform statistics on local mutation densities of bins based on genomic coverage, we calculated the mean coverage of each bin and performed linear regression between mutation densities and coverage for each sample displaying the mean and standard deviation and regression line on the graph. To generate mutation density profiles across regions, windows were generated within, upstream or downstream, each site of the region separately according to the number of bins and number of bases flank specified. Where regions contain sites of varied lengths e.g. gene bodies, the number of windows for each site was fixed therefore changing the number of bases per window in the region. Mutation densities were then calculated in each of the windows as described above.

### Mutation Trinucleotide Frequency Calculations

Mutation trinucleotide frequencies were calculated using DeconstructSigs [36]. For frequencies across the whole genome, ‘genome’ normalisation was used. For frequencies on specific regions, such as CBS motifs, the trinucleotide composition was calculated using grep scripts and then these were input into DeconstructSigs with ‘manual’ normalisation.

### Genomic Annotations and Data Binning

Gene expression data was taken from GTEx portal and the top half of expressed genes were defined as ‘expressed’ in bladder. Genes with 0 counts in bladder were defined as ‘silent’. Annotations of genic regions including 5’ untranslated region (UTR), 3’ UTR, exons and introns were accessed from UCSC table browser for hg19. Intergenic regions for hg19 were defined as parts of the genome without overlap of any of these regions. Hg19 coverage and narrow peaks data for human bladder tissue DNase-seq experiments were accessed from ENCODE [37] (ENCSR813CKU) (Supplementary table 1) as bigWig and bed file respectively. ChIP-seq for XPD, XPB and input was accessed from GEO (GSE44849) and cisplatin/ oxoplatin based TFIIH XR-seq was accessed from GEO under accession GSE82213 as bigwig files . Deeptools ‘bigWigCompare’ was used to generate ChIP to input log2 ratio bedgraph files for XPD and XPB ChIP-seq. For TFIIH XR-seq data, ‘bigWigMerge’ was used to merge plus and minus outputting a bedgraph. 1 kb windows of hg19 were generated and then filtered for blacklisted and low coverage regions of the genome. To divide the genome into bins based on coverage of different genomic assays, including DNase-seq, replication time, XPD ChIP-seq and TFIIH, the mean bedgraph signal from genomic assays was calculated for each of the 1kb filtered genomic windows using bedtools map [35]. For mutation density calculations, these filtered 1 kb windows were then divided into quintiles based on coverage from lowest signal (bin 1) to highest signal (bin 5).

Bladder DHS peaks were overlapped with other DHS marks to generate annotations for bladder DNase hypersensitive regions (DHS) as follows. Promoters were defined by overlap with bladder H3K4Me3 ChIP-seq peaks from ENCODE (ENCSR632OWD) (Supplementary table 1) and then gene start sites to get promoters. Bladder DHS peaks were overlapped with high quality, experimentally determined CBS accessed from supplementary materials of [16] to generate CBS annotations. Later analysis of CBS uses these high quality CBS annotations [16] without overlapping with bladder DHS. Finally, enhancers were defined as the centre of bladder H3K27Ac ChIP-seq peaks from ENCODE (ENCSR054BKO) (Supplementary table 1) that overlapped bladder DHS peaks. Chromatin A/B compartments for bladder were taken from supplementary files of [38]. For later analysis on CBS, all 31252 CBS sites were used [16].

The above annotations were converted to hg38 using ‘liftOver’.

### Generating Coverage Profiles Across Genomic Regions for ChIP-seq Data

BigWig files for XPD and XPB ChIP-seq and input were accessed from gene expression omnibus (GEO) under accession GSE44849 which was previously published [39] (Supplementary table 1). Deeptools ‘bigwigCompare’ [40] was used to generate log2 ratio bigwig files of the ChIP compared with input, skipping regions that had no coverage in both the input and the ChIP. To generate sequencing coverage profile plots for regions, windows were generated within, upstream or downstream of each site in the region according to the number of bins and number of bases flank specified. Where regions contain sites of varied lengths e.g. gene bodies, the number of windows for each site was fixed therefore changing the number of bases per window. UCSC tool ‘bigWigAverageOverBed’ was then used to retrieve the average signal of each region in each window. The average region signal of each window was averaged and plotted positionally.

### Uracil Sequencing Data

Uracil sequencing data previously published [41] was accessed from European Nucleotide Archive under accession PRJNA728500. Data was processed as the authors described. Briefly, fastq files were trimmed with bbduk and aligned to hg19 using bwa [42], then duplicates were removed using picard (http://broadinstitute.github.io/picard). Single base locations of uracil were located using authors scripts “fetch_dU_by_chrom.py” from https://github.com/Jyyin333/Ucaps-seq. If the reference base matched a T/A or a C/G it was considered to be from incorporation or deamination respectively. Uracil sequencing data profiles were drawn around hg19 CBS sites using coverageBed [35]. Bedtools slop and fastaFromBed were used to retrieve the trinucleotide sequence context of uracil.

### Support Machine Vector Model

Results of somatic mutation densities from GE for the following genomic regions: CBS motif, 1000 bp flanking CBS motif, coding exons, introns, 3’UTR, 5’UTR, intergenic, open and closed chromatin and 5 regions of the genome binned by replication time, DNase hypersensitivity, XPD ChIP-seq coverage and TFIIH XR-seq coverage was used as the input to train a support vector machine (SVM) for classifying whether a sample was a driver or passenger mutation. The svm function from the e1071 R package was used. The SVM model was first evaluated using leave-one-out-cross validation using the GE cohort (samples with well defined ERCC2 mutation status). We further trained the SVM model using all of these GE samples and tested this on the independent TCGA cohort. Finally, we used the same model to evaluate the ERCC2 mutation status of the GE samples with undefined ERCC2 mutation status.

## Data Availability

Accessions and references to dataset used in the paper are listed in the Supplementary Table 1. Those with access to Genomics England Research Environment may request the locations of all scripts required to complete analyses.

## Contributions

J.A.B. analysed data, prepared figures, and wrote the manuscript; H.F., N.C.Y. X.Z., H.Y. Y.T.W. analysed and interpreted data, T.O., M.W.W, N.A.B. contributed data, S.W. contributed data and jointly supervised the research; J.W.H.W. conceived and designed the study, analysed data, revised manuscript and supervised the research.

## Acknowledgements

This work was enabled by access to data and findings generated by the 100,000 Genomes Project, under the Pan-Cancer Genomics England Clinical Interpretation Partnership (GeCIP), project ID 752. The project was supported by the Research Grants Council, HK (17100920, R7022-20 and C7028-19G) to JWHW and the Centre for Oncology and Immunology under the Health@InnoHK Initiative funded by the Innovation and Technology Commission, The Government of Hong Kong SAR, China.

## Competing interests

The authors have declared that no competing interests exist.

## Supplementary Figures

**Figure S1.**
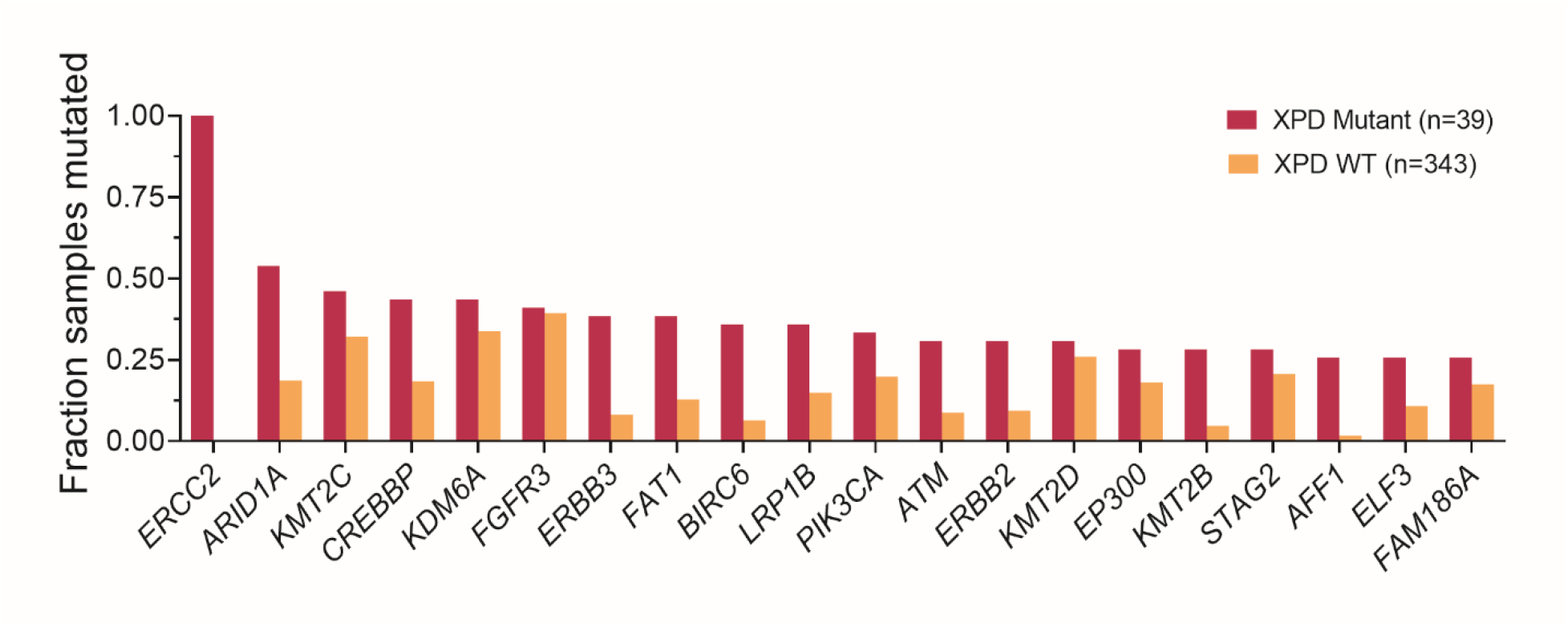
Contribution of different mutations in XPD mutant and wild-type (WT) BLCA. The fraction of XPD mutant and wild-type (WT) samples with protein-altering mutations (missense, nonsense, frameshift) in oncogenes defined from IntOGen database. The most frequently occurring 20 genes are shown.

**Figure S2.**
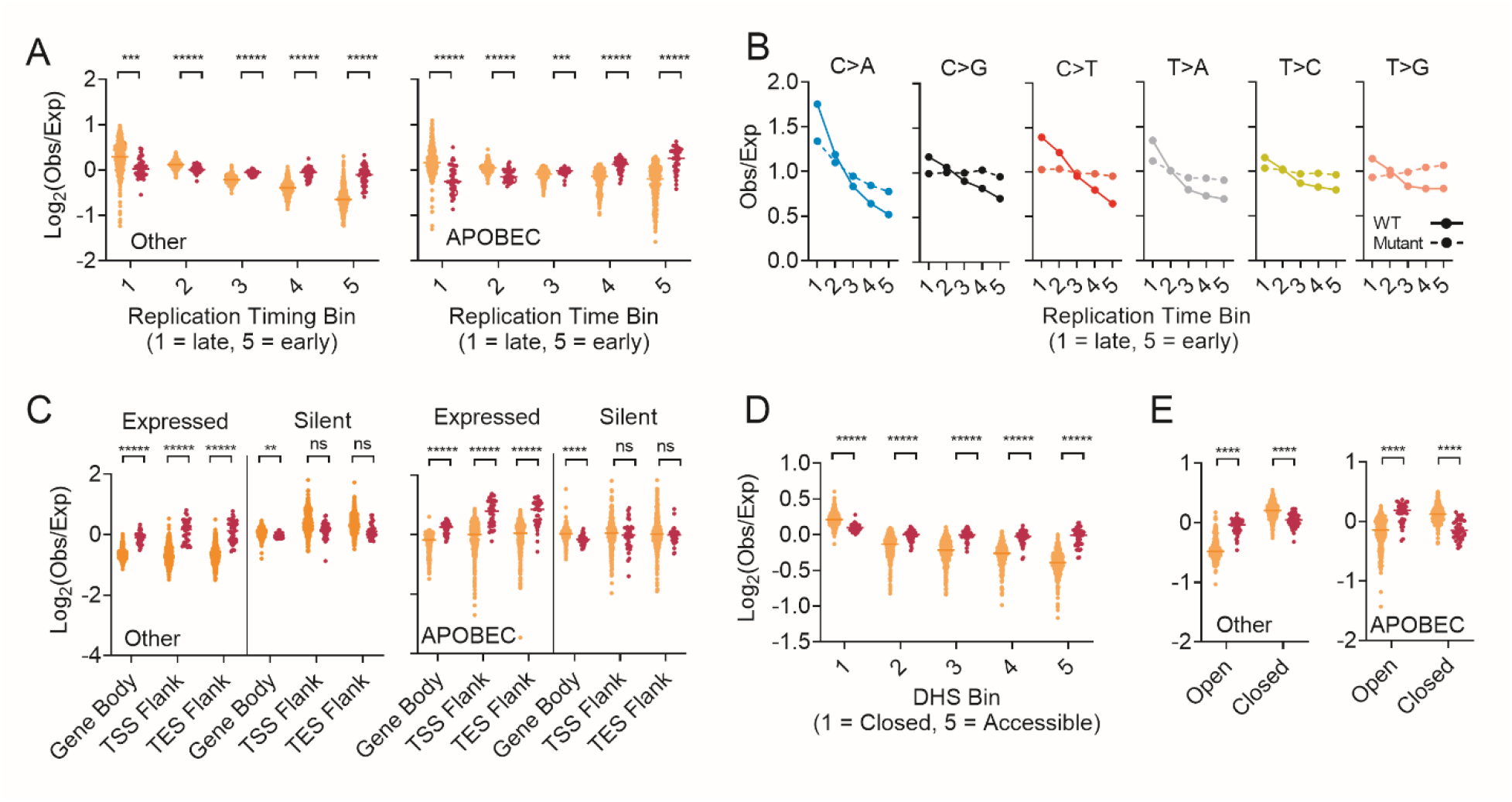
Genome wide distribution of mutations in XPD Mutant and WT BLCA. (A) Mutation densities of individual data points with respect to replication time bins for Other and APOBEC SNVs. (B) Relationship between Other mutations and replication time separated for C>A, C>G, C.T, T>A, T>C and T>G (C) A statistical representation of plots in Fig 2C. Mutation densities of individual samples are shown for the gene body, the 2.5 Kb upstream flanking region of TSS (TSS flank) and 2.5 Kb downstream flanking region of TES (TES flank). (D) and (E) Analysis in figure 2D and 2E performed with CBS and genes subtracted from DHS bins and from open and closed A/B compartments. *** q < 0.0001, **** q < 0.00001, ***** q < 0.000001, n.s. not significant, Student’s t-test with multiple testing correction.

**Figure S3.**
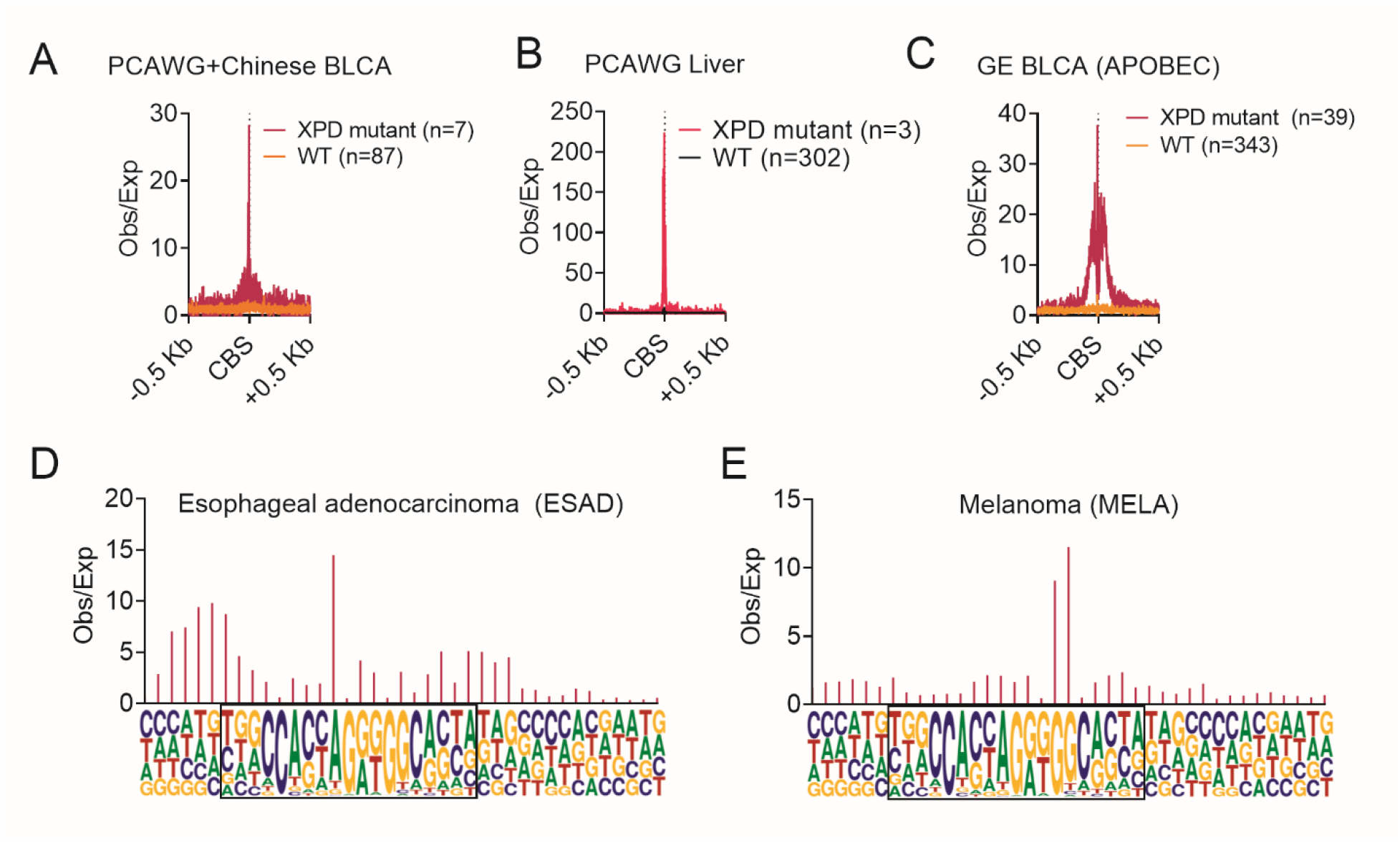
Mutation profiles across CBS and flanking regions. Mutation profiles across CBS of for samples from 3 additional cohorts of BLCA (A), PCAWG liver cancer (B) and APOBEC SNVs for WT and XPD mutant GE BLCA (C). Observed/Expected mutation profile across the CTCF motif for esophageal adenomcarcinoma (ESAD) (D) and melanoma (MELA) (E).

**Figure S4.**
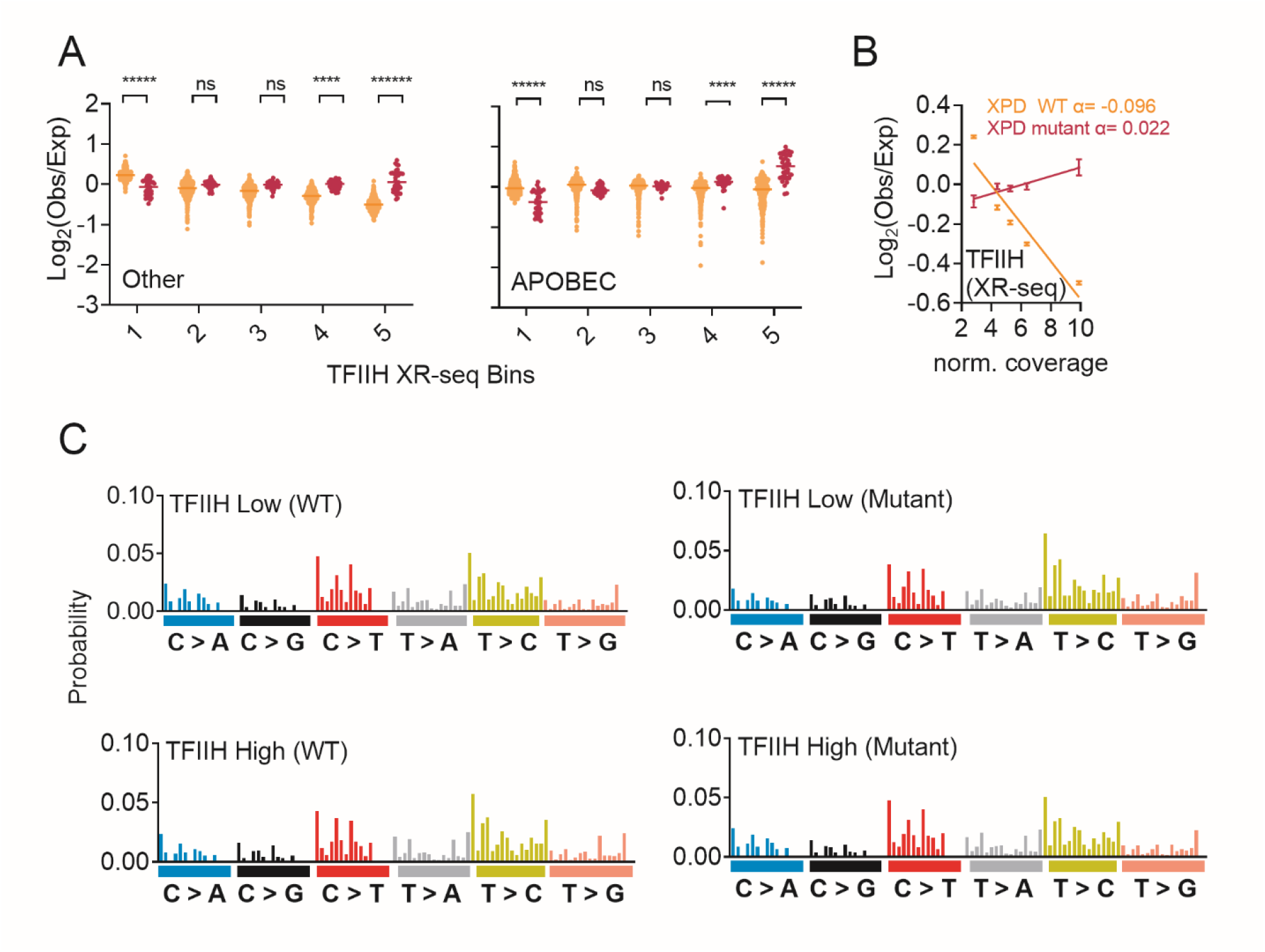
Mutation load and mutational signature in relation to XPD binding and TFIIH XR-seq coverage. (A) Observed/Expected mutation load for Other and APOBEC SNVs across TFIIH XR-seq coverage bins for Other SNVs. *** q < 0.0001, **** q < 0.00001, ***** q < 0.000001, n.s. not significant, Student’s t-test with multiple testing correction. (B) regression of S4A for BLCA (left) and ESAD (right) (C) Trinucleotide spectrum of mutations in high and low TFIIH XR-seq regions in WT (left) and XPD mutant (right) GE BLCA.

**Figure S5.**
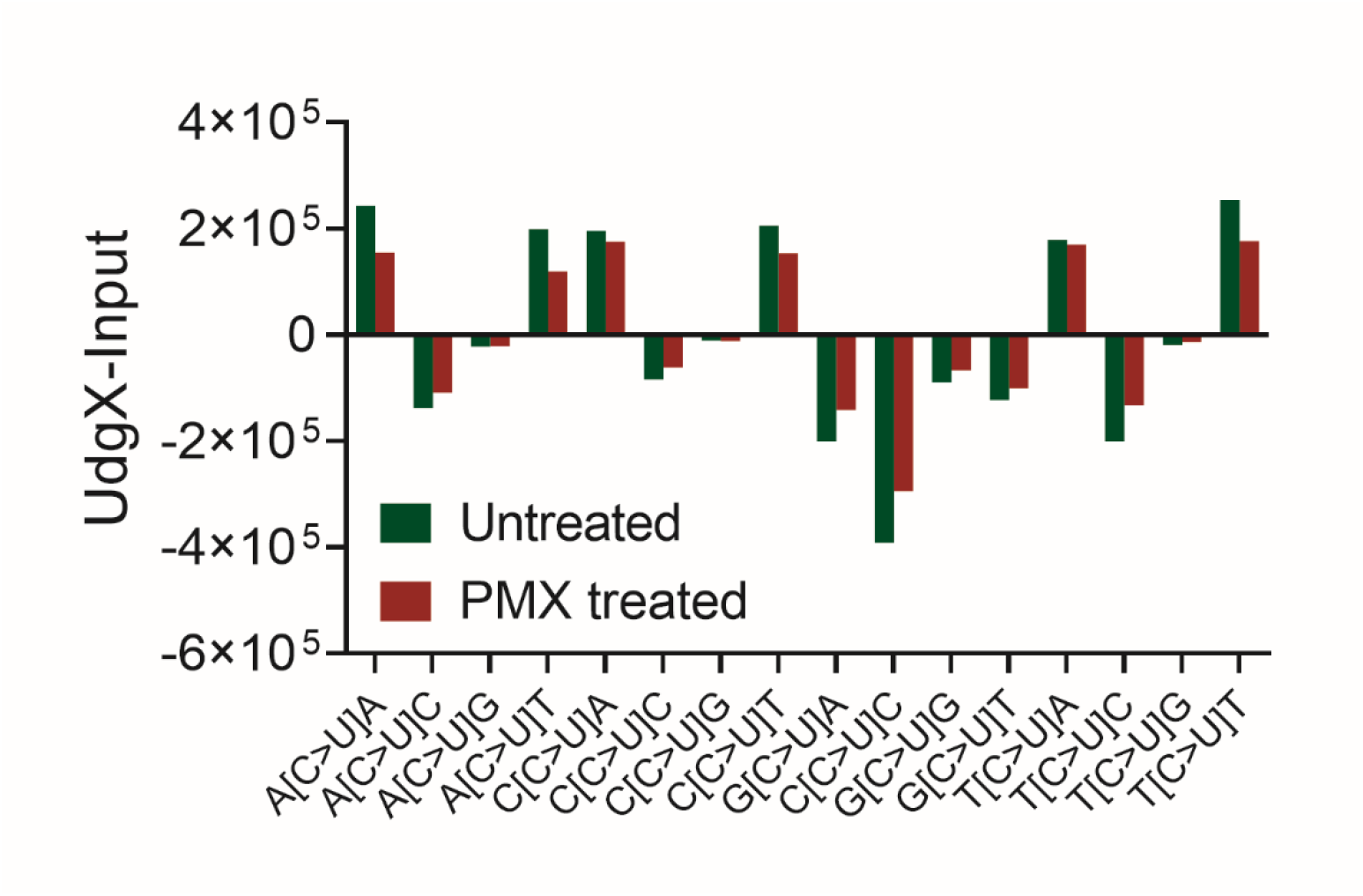
Trincleotide context of genomic uracil sites associated with cytidine deamination. Frequency of single base sites of uracil associated with cytosine deamination in the trinucleotide context as UdgX sequencing exp-input. C>U indicates a uracil was detected where the reference base is a cytosine.

## Supplementary Tables

**Supplementary table 1.**
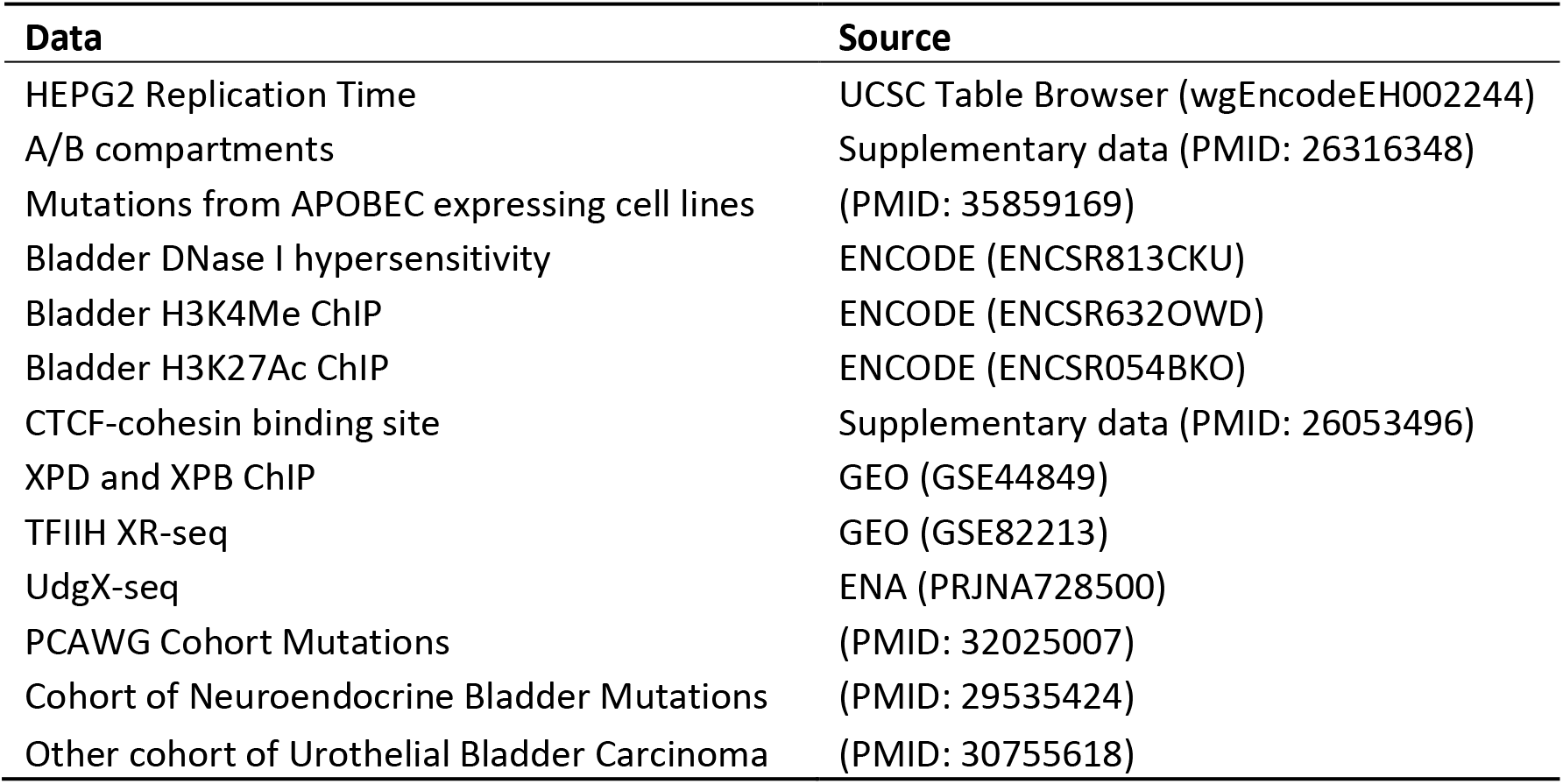
List of public datasets used in the study.

| **Data** | **Source** |
| --- | --- |
| HEPG2 Replication Time | UCSC Table Browser (wgEncodeEH002244) |
| A/B compartments | Supplementary data (PMID: 26316348) |
| Mutations from APOBEC expressing cell lines | (PMID: 35859169) |
| Bladder DNase I hypersensitivity | ENCODE (ENCSR813CKU) |
| Bladder H3K4Me ChIP | ENCODE (ENCSR632OWD) |
| Bladder H3K27Ac ChIP | ENCODE (ENCSR054BKO) |
| CTCF-cohesin binding site | Supplementary data (PMID: 26053496) |
| XPD and XPB ChIP | GEO (GSE44849) |
| TFIIH XR-seq | GEO (GSE82213) |
| UdgX-seq | ENA (PRJNA728500) |
| PCAWG Cohort Mutations | (PMID: 32025007) |
| Cohort of Neuroendocrine Bladder Mutations | (PMID: 29535424) |
| Other cohort of Urothelial Bladder Carcinoma | (PMID: 30755618) |

**Supplementary table 2.** XPD Mutations and pathogenicity classification in cohort.

| **Mutation** | **Pathogenicity** | **Number of samples** |
| --- | --- | --- |
| Arg487T | ERCC2_hotspot | 1 |
| Asn238Ser | ERCC2_hotspot | 6 |
| Gln662His | ERCC2_not_hotspot | 1 |
| Glu606Gln | ERCC2_hotspot | 4 |
| Glu606Lys | ERCC2_hotspot | 2 |
| Glu79Ile80Phe | ERCC2_hotspot | 1 |
| Glu86A | ERCC2_hotspot | 1 |
| Glu86Val | ERCC2_not_hotspot | 1 |
| Gly607Ala | ERCC2_not_hotspot | 1 |
| Gly665Ala | ERCC2_not_hotspot | 1 |
| Gly665Ser | ERCC2_hotspot | 1 |
| Gly675Cys | ERCC2_hotspot | 1 |
| Gly675Ser;Ser74Leu | ERCC2_hotspot | 1 |
| Gly675Val | ERCC2_hotspot | 1 |
| His237Gln | ERCC2_hotspot | 1 |
| His659Arg | ERCC2_not_hotspot | 1 |
| Phe157Leu | ERCC2_not_hotspot | 1 |
| Pro463Ala | ERCC2_not_hotspot | 1 |
| Pro463Gln;Ser44Leu | ERCC2_hotspot | 1 |
| Ser44Leu | ERCC2_hotspot | 1 |
| Ser74Pro | ERCC2_hotspot | 1 |
| splice_region_variant | ERCC2_not_hotspot | 1 |
| splice_region_variant;Phe651Val | ERCC2_not_hotspot | 1 |
| Thr460Ser | ERCC2_not_hotspot | 1 |
| Thr484Met | ERCC2_hotspot | 5 |
| Tyr14Cys | ERCC2_hotspot | 2 |
| Tyr24Cys | ERCC2_hotspot | 3 |
| Tyr72Cys | ERCC2_hotspot | 5 |
| Val242Phe | ERCC2_hotspot | 1 |

## Notes

### Competing Interest Statement

The authors have declared no competing interest.

### Summary of Updates

The analysis has been updated based on data from the Genomics England cohort. All figures in the manuscript has been updated although major conclusion has not changed.

